# Brain BOLD MRI O_2_ and CO_2_ stress testing: Implications for perioperative neurocognitive disorder following surgery

**DOI:** 10.1101/619361

**Authors:** W. Alan C. Mutch, Renée El-Gabalawy, Lawrence Ryner, Josep Puig, Marco Essig, Kayla Kilborn, Kelsi Fidler, M. Ruth Graham

## Abstract

Respiratory end-tidal (ET) gas control is fundamental to anaesthetic management. The range of ET O_2_ and CO_2_ during the conduct of anaesthesia can significantly deviate from values in the awake state. Recent work shows ET CO_2_ influences the incidence of perioperative neurocognitive disorder (POND). We examine the effects of controlled alterations in both ET O_2_ and CO_2_ on cerebral blood flow (CBF) in awake adults using BOLD MRI. Twelve healthy adults had BOLD and CBF responses measured to alterations in ET CO_2_ and O_2_ in various combinations commonly observed under anaesthesia. Dynamic alterations in regional BOLD and CBF were seen in all subjects with expected and inverse responses to both stimuli. These effects were incremental and rapid (within seconds). The most dramatic effects were seen with combined hyperoxia and hypocapnia. Inverse responses increased with age. Here we show that human brain CBF responds dramatically to alterations in ET respiratory gas tensions commonly seen during anaesthesia. Such alterations may impact the observed incidence of POND following surgery and intensive care, and is an important area for further investigation.

---

Perioperative neurocognitive disorder (POND) is a newly proposed nomenclature to classify cognitive problems that can manifest after anaesthesia and surgery.^1^ The term encompasses pre-existing problems such as mild cognitive impairment or dementia, acute postoperative delirium, and postoperative cognitive dysfunction, with the potential for learning and memory deficits in young children and a progressive downward arc in cognitive abilities in older adults after surgery. Despite efforts to apply consistent terminology to facilitate research in this area, the phenomenon of POND continues to be poorly understood. The cost to society of this problem is immense and new insights are desperately needed.^234^ Putative causative factors include, first, the potential neurotoxicity of the anaesthetic agents themselves ^5,6,7,8^. The basic premise is that the neurotransmitter modulating effects of anaesthetic agents damage neurons or their dendrites at the extremes of life and lead to caspase induced apotosis.^9,10^ Evidence supporting this is unequivocal in animal models up to and including non-human primates.^11,12^ However, translation to the human domain is contradictory and an increasing series of clinical studies, both in children and adults have failed to provide the expected surrogates for evidence of neuronal damage as seen in animal models of anaesthetic neurotoxicity. Many of the animal models are highly constrained by specific requirements to demonstrate neuronal injury. An overview of some of these translational issues has been recently highlighted.^13,14,15^ The large retrospective studies that have failed to show an association between anaesthetic exposure or multiple exposures and neurocognitive deficits may be criticized as not being fine grained enough to address the problem. However, more recent prospective trials also fail to show a signal of POND either in children or adults. For example, a paediatric study failed to show any difference in the primary outcome of individual intelligence quotient testing later in childhood when anaesthesia naive children were compared to propensity matched children undergoing surgery at less than 3 years of age with either single or multiple anaesthetic exposure.^16^ An adult study failed to show a decrease in postoperative delirium in older adults with anaesthetic depth minimized by limiting EEG burst suppression compared to those managed conventionally with a greater depth of anaesthesia.^17^ In light of these null findings, new insights appear warranted.

Alternative considerations include neuroinflammatory effects consequent with surgical stress^18,19,20^, and premorbid risk factors (e.g., cognitive dysfunction, psychiatric illness, increasing age) which may influence the incidence of POND.^3,21,22^ More recent work suggests that a multi-factorial approach, described as a stress-diathesis model^23^, may provide further insights. In this model, the conduct of anaesthesia, which includes the management of respiratory gases during mechanical ventilation, may contribute to the stress of the perioperative period in vulnerable individuals (the diathesis) and is associated with development of POND.^23^

Although respiratory end-tidal gas control is a fundamental principle of anaesthetic management, the range of end-tidal (ET) O_2_ and CO_2_ typically seen during the conduct of anaesthesia can significantly deviate from the awake baseline state, but has not been considered as a potential contributor to POND in previous investigations. Periods of both hypocapnia during mechanical ventilation and hypercapnia during the reestablishment of spontaneous ventilation are common. The use of 100% oxygen during anaesthesia induction and emergence and maintenance of anaesthesia with O_2_ concentrations of 50% are typical.

The effects of these alterations in respiratory gases have significant effects on cerebral blood flow and oxygenation which may be implicated in cognitive dysfunction. In the present study, using blood oxygenation level dependent (BOLD) and pseudo continuous arterial spin labeling (pCASL) MRI, we examine the effects of controlled alterations in ET O_2_ and CO_2_ in various combinations commonly observed during anaesthesia and surgery on cerebral blood flow (CBF) in 12 healthy awake adults. Based on the significant alterations in CBF observed, we propose mechanisms to suggest that manipulation of the gases of respiration (O_2_ and CO_2_) may be a significant contributor to the intraoperative stress leading to POND in vulnerable individuals.^24,25^ The proposed mechanisms described here can be easily examined and ruled in or out based on well-conducted clinical trials.

## Methods

These studies relate to ongoing examination of volunteer subjects and patients undergoing surgical intervention and concussion studies at the University of Manitoba (U of M), Winnipeg, Canada. The protocol was approved by the Biomedical Research Ethics Board (BREB), at the U of M, and a cohort of the study group was registered at ClincialTrials.gov for some components of the study NCT02126215. For the present study, 12 healthy adult volunteers gave witnessed informed consent and underwent MRI while ET gases were manipulated in a controlled fashion with a RespirAct (a computer-controlled model-based prospective end-tidal gas mixer).^26,27^ The volunteers had no neurological health issues nor were receiving psychotropic drugs for any psychiatric conditions. None had a history of POND with any previous surgery.

### End-tidal gas protocols

Following a brief overview, subjects had an air-tight plastic mask affixed to their face. They then were positioned on the imaging table and a prep sequence was run on the RespirAct to determine the individual’s resting ET gases, breathing frequency and tidal volume. When determined, the subject was positioned in the magnet bore. Breathing sequences (see Supplemental File 1 for an example) included i) a CO_2_ ramp protocol; an 11-minute sequence of baseline ET CO_2_ (1-min), a square wave increase in ET CO_2_ by 5 mm Hg (1-min), a return to baseline (1-min), a square wave period of spontaneous hyperventilation upon command to decrease ET CO_2_ by 5 mm Hg (1-min), a 6-min ramp increase in CO_2_ to 10 mm Hg above baseline, a 1-min return to baseline CO_2_. During this sequence the ET O_2_ was clamped at 100 mm Hg; ii) an O_2_ ramp protocol; an 11-minute sequence of baseline ET O_2_ fixed at 100 mm Hg (1-min), a square wave increase in ET O_2_ to 500 mm Hg (1-min, 400 mm Hg above baseline), a return to baseline (2-min), a 6-min ramp increase in O_2_ to 400 mm Hg above baseline, a 1-min return to baseline O_2_. During this sequence the ET CO_2_ was clamped at the subject’s baseline determined from the initial CO_2_ ramp sequence; iii) pCASL sequence 1 (a 3-min baseline sequence and a block design square wave alteration); baseline O_2_ for 3-min, then square wave increase in O_2_ to 400 mm Hg (hyperoxia) with ET CO_2_ clamped at subject baseline (normocapnia): iv) pCASL sequence 2, after a 5-minute re-equilibration period; baseline CO_2_ for 3-min, then square wave increase in CO_2_ to 5 mm Hg above baseline (hypercapnia), with ET O_2_ clamped at 100 mm Hg (normoxia): pCASL sequence 3, after a 5-minute re-equilibration period; baseline CO_2_ and ET O_2_ at 100 mm Hg for 3-min, then square wave decrease in CO_2_ by 5 mm Hg below baseline (hypocapnia) by spontaneous hyperventilation on command and ET O_2_ increased to 400 mm Hg (hyperoxia).

### Imaging protocols

Imaging occurred on a Siemens 3T Verio magnet. The head coil had 12-channels. Each subject underwent gradient field mapping, magnetization-prepared rapid acquisition with gradient echo (MPRAGE) anatomic imaging, gradient recalled echo (GRE) BOLD imaging × 2 for the ramp protocols, an M0 calibration image for each of the pCASL sequences, pCASL × 3 for cerebral blood flow determination during the various end-tidal gas protocols, axial fluid attenuated inversion recovery (AX-FLAIR) and AX-T2* GRE. The BOLD GRE images were obtained using Siemens proprietary software for prospective motion correction (PACE). See Supplemental File 2 for a fuller description of the imaging sequences. Total imaging time was approximately 45-60 minutes/session.

### Post-processing of images

The images were converted to .nii files and using custom written batch files preprocessing was accomplished using standard SPM8 pipelines. BOLD images were re-aligned, co-registered to the MPRAGE images, normalized and smoothed with a 5×5×5 mm smoothing kernel at FWHM. These images were then fit to a GLM based on the end-tidal CO_2_ and O_2_ ramps respectively. Both the expected and the inverse responses were examined at various p-values. First and 2^nd^ level analysis were undertaken for the images; the 2^nd^ level using a leave-one-out approach. Various masks were applied to the images to examine specific regions and control for excessive alteration in B0 field strength. See Supplemental File 3 for a description of the masks used. The pCASL images were processed using the ASLtbx developed by Ze Wang.^28^ Cerebral blood flow as determined at the baseline settings for each of the three runs and the altered CBF determined for cerebrovascular reactivity (CVR) assessment of the block-designed change in flow. Flow maps were determined for baseline settings, the block-designed stress and the difference between.

## Results

Twelve studies were undertaken. In all, CO_2_ and O_2_ BOLD GRE ramp protocols were undertaken and completed. In 6, pCASL measurements were obtained. In these, the first two pCASL protocols were completed in all cases and all 3 pCASL protocols were completed in 4 subjects. There were no complications noted following completion of the protocols.

Subject demographics are seen in Table 1. One subject had a history of migraine headaches and one subject had a prior history (remote) of concussion.

**Table 1.** Subject Demographics.

| Study # | Age | Sex | Status | Medications | Resting BP |
| --- | --- | --- | --- | --- | --- |
| 1 | 42 | M | Migraines | Nil | 122/77 |
| 2 | 29 | M | Remote concussion | Nil | 118/69 |
| 3 | 61 | F | Healthy | Antibiotics | 135/83 |
| 4 | 45 | M | Healthy | Nil | 138/86 |
| 5 | 57 | F | Healthy | Nil | 141/89 |
| 6 | 53 | F | Healthy | Nil | 145/77 |
| 7 | 62 | M | Healthy | Nil | 155/84 |
| 8 | 44 | M | Healthy | Nil | 110/70 |
| 9 | 20 | F | Healthy | Nil | 110/66 |
| 10 | 54 | F | Healthy | Nil | 130/82 |
| 11 | 56 | F | Healthy | Nil | 138/74 |
| 12 | 52 | F | Healthy | Nil | 112/65 |

The end-tidal gas results are shown in Table 2. All subjects had ET gas control close to their planned alterations. A plateau was seen for the upper limit of ET O_2_ for all with a maximum in the range of 475 mm Hg in the O_2_ ramp protocol. The 1^st^ level analysis from the SPM processing fit to the general linear model (GLM) for the CO_2_ ramp is shown in Table 3. The raw voxel counts are with an individualised grey and white matter inclusive mask only and the masked voxel counts are following application of the individualised mask as shown in Supplemental File 3. The 1^st^ level analysis from the SPM processing fit to the general linear model (GLM) for the O_2_ ramp is shown in Table 4. Table 5 and 6 show the 2^nd^ level analysis for CO_2_ and O_2_ ramps respectively. A group comparison of the 4 individuals with masked voxels counts greater than 50 for inverse responses to the CO_2_ and O_2_ ramps is shown in Supplemental File 4. An example of 1 subject for the brain BOLD and inverse response to the CO_2_ ramp at clamped O_2_ is shown in Figure 1 and the BOLD response in the same patient to the O_2_ ramp at clamped CO_2_ is shown in Figure 2. There is an association with increasing inverse voxel counts response to CO_2_ and O_2_ with increasing age. This correlation best fit a 2^nd^ order polynomial; r = 0.664, p = 0.019 (Figure 3). A result from the 2^nd^ level analysis is also shown. An individual response to the pCASL protocol is shown in Figure 4. For this subject the pCASL CVR responses are shown for hypercapnia at isoxia (A), hyperoxia at isocapnia (B) and hyperoxia with hypocapnia (C). With hypocapnia the decrement in CBF becomes marked. The heterogeneity of BOLD to CBF response to the O_2_ ramp and hyperoxia-baseline CVR response is shown in Figure 5. The decrease in CBF with decreased BOLD signal is highlighted for the frontal cortex. The inverse response to hyperoxia is associated with a regional decrement in CBF. The dynamism of responsiveness to alterations in respiratory gases is evident.

**Figure 1 A-C:**
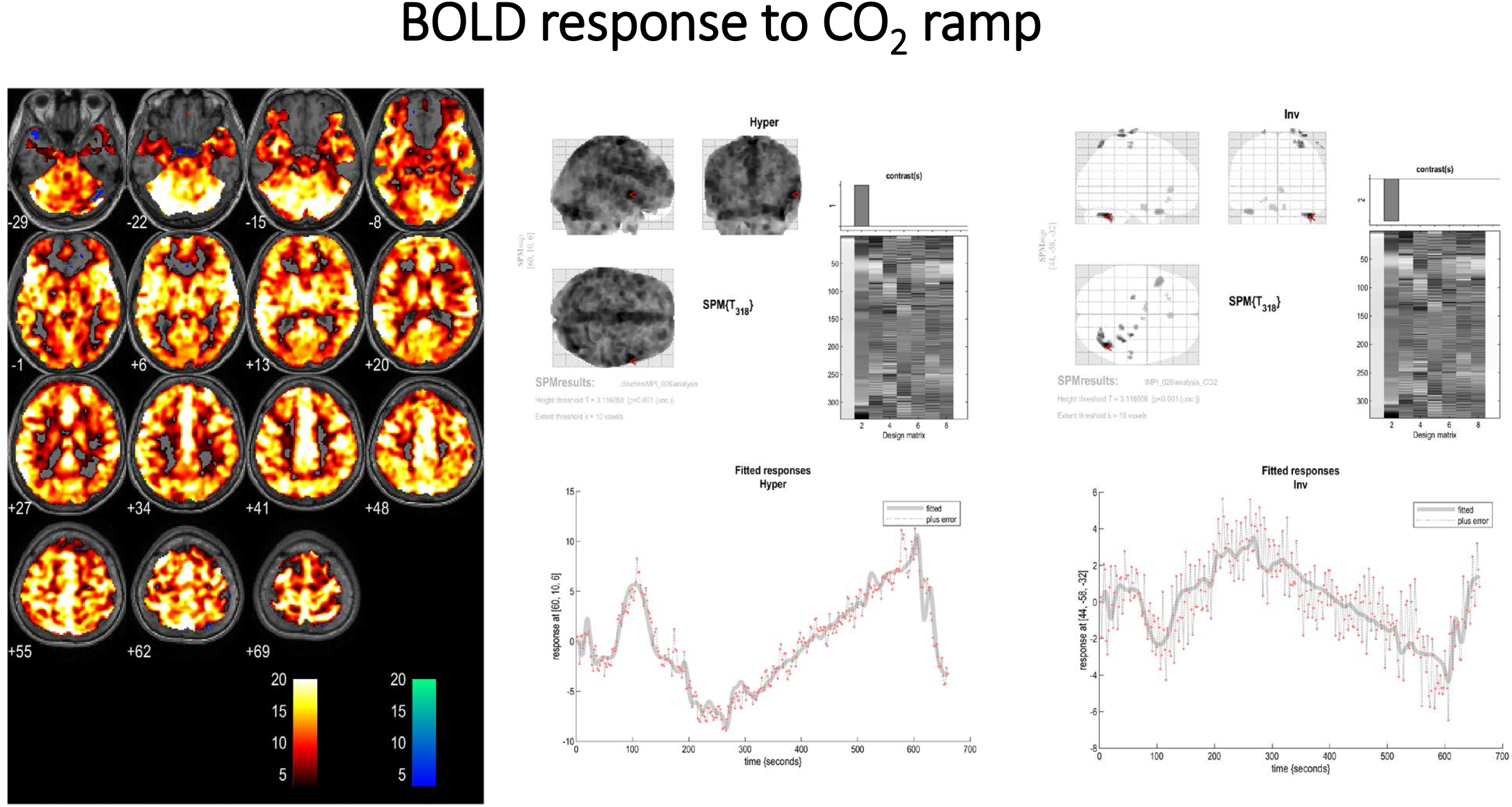
A: BOLD response to the CO_2_ ramp with the ETO_2_ tension clamped at ~100 mmHg. The expected response with increased BOLD signal to the ramp stimulus is depicted by orange voxels. A diffuse response is seen. The scale is t-scores based on fit to the GLM from SPM first level analysis. The blue voxels depict the inverse response. The t-score had to exceed 3.11 (p=0.001) to be colourized. B: The response of one voxel to the CO_2_ ramp. The incremental and rapid increase with the step change in CO_2_ (grey scale) is seen by the red dots representing the BOLD scan signal intensity at that moment in time. C: The inverse response of one voxel to the CO_2_ ramp. The incremental and rapid decrease with the step change in CO_2_ (grey scale) is seen by the red dots representing the BOLD scan signal intensity at that moment in time.

**Figure 2 A-C:**
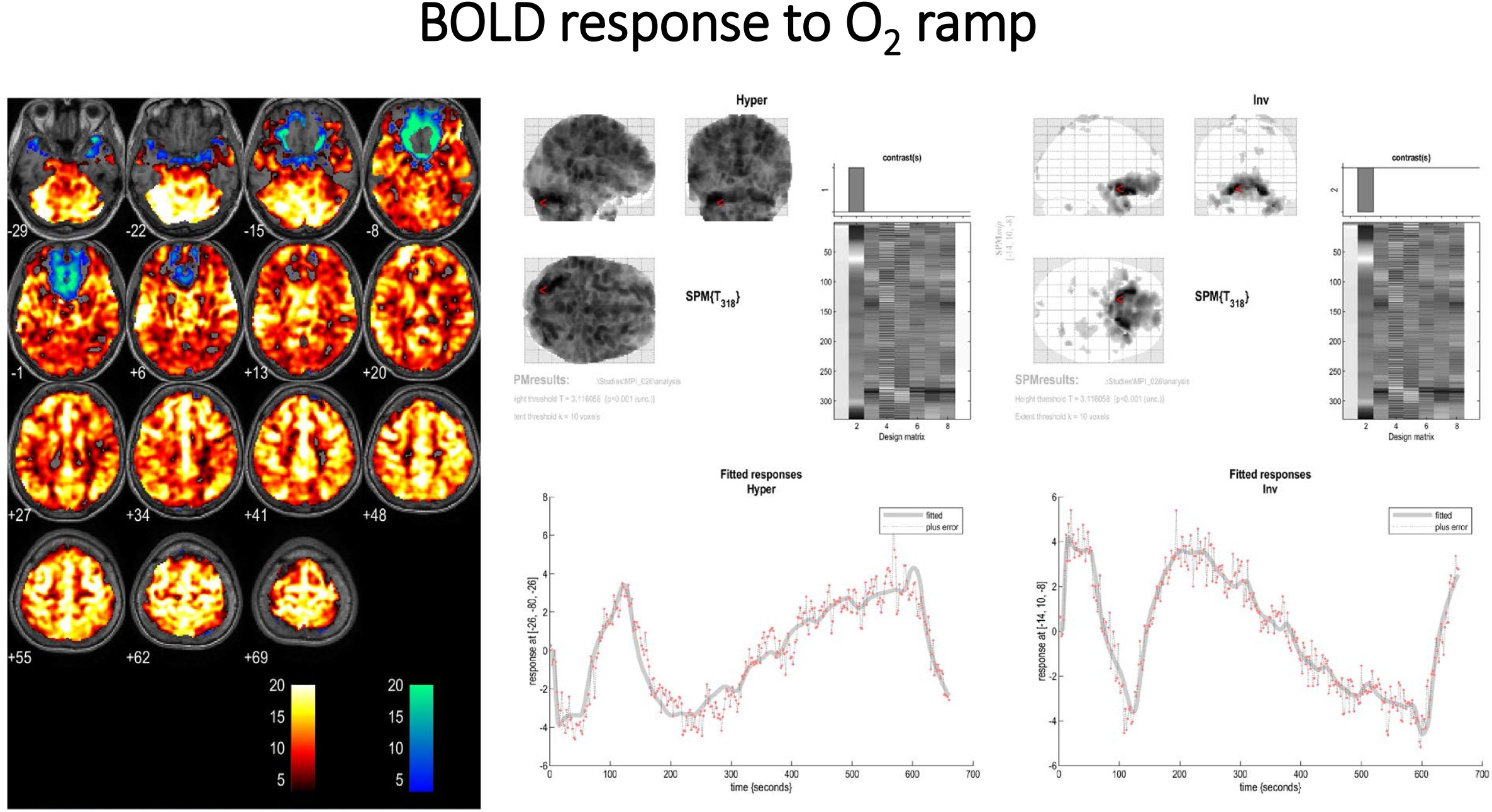
BOLD response to the O_2_ ramp with the ETCO_2_ tension clamped at ~43 mmHg. The expected response to the ramp stimulus is depicted by orange voxels. A diffuse response is seen. The scale is t-scores based on fit to the GLM from SPM first level analysis. The blue voxels depict the inverse response. The t-score had to exceed 3.11 (p=0.001) to be colourized. B: The response of one voxel to the O_2_ ramp. The incremental and rapid increase with the step change in O_2_ (grey scale) is seen by the red dots representing the BOLD scan signal intensity at that moment in time. C: The inverse response of one voxel to the O_2_ ramp. The incremental and rapid decrease with the step change in O_2_ (grey scale) is seen by the red dots representing the BOLD scan signal intensity at that moment in time.

**Figure 3:**
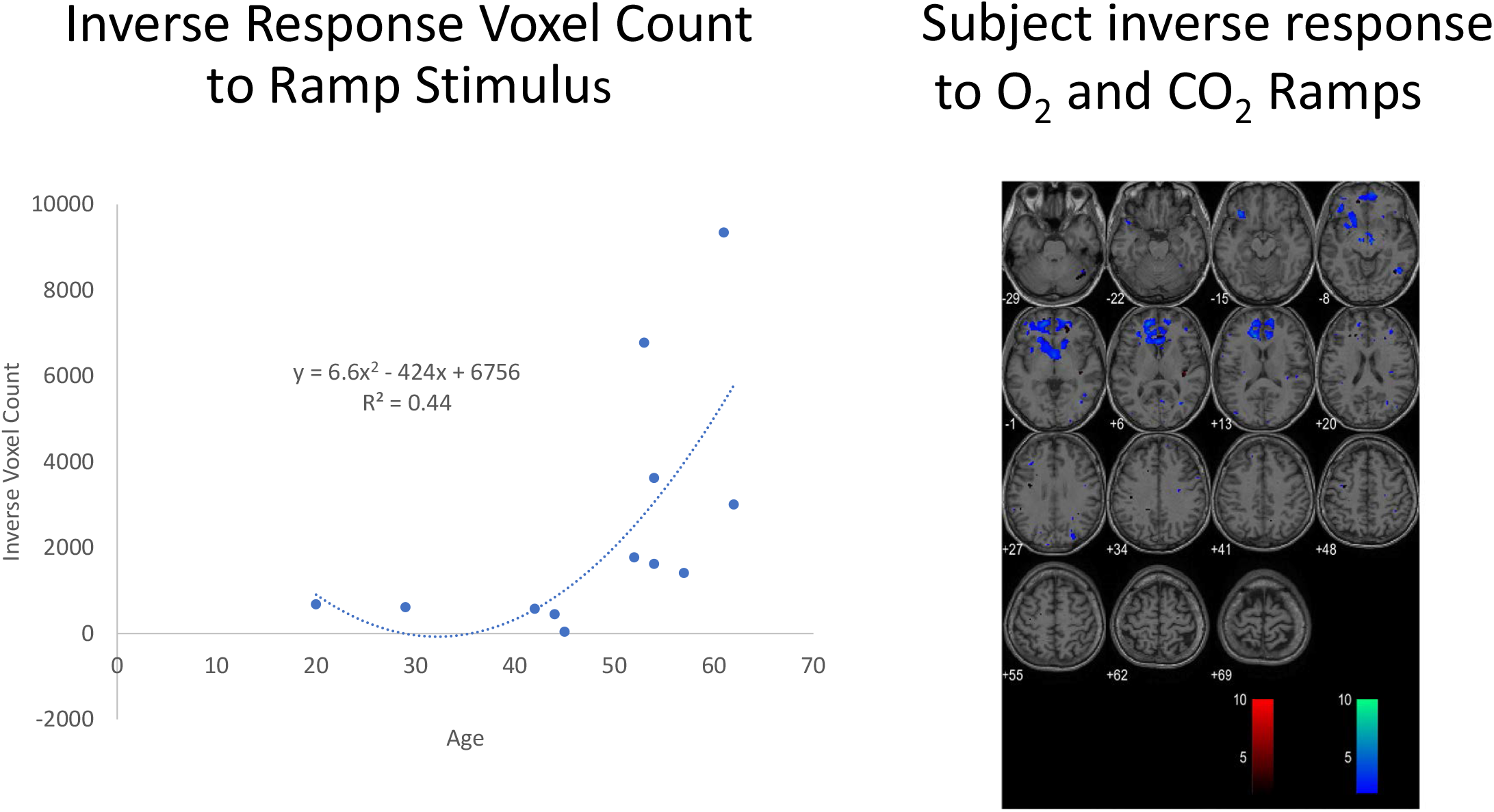
Graph of the relationship between inverse voxels to CO_2_ and O_2_ and age. The data best fit a 2^nd^ order polynomial; p=0.019. A representative example in one patient following 2^nd^ level analysis showing the less than response to the CO_2_ ramp stimulus (in red) and the less than response to the O_2_ ramp stimulus (in blue) with a leave-one-out analysis (removing this subject and comparing this subject to the other 11 subjects).

**Figure 4 A-C:**
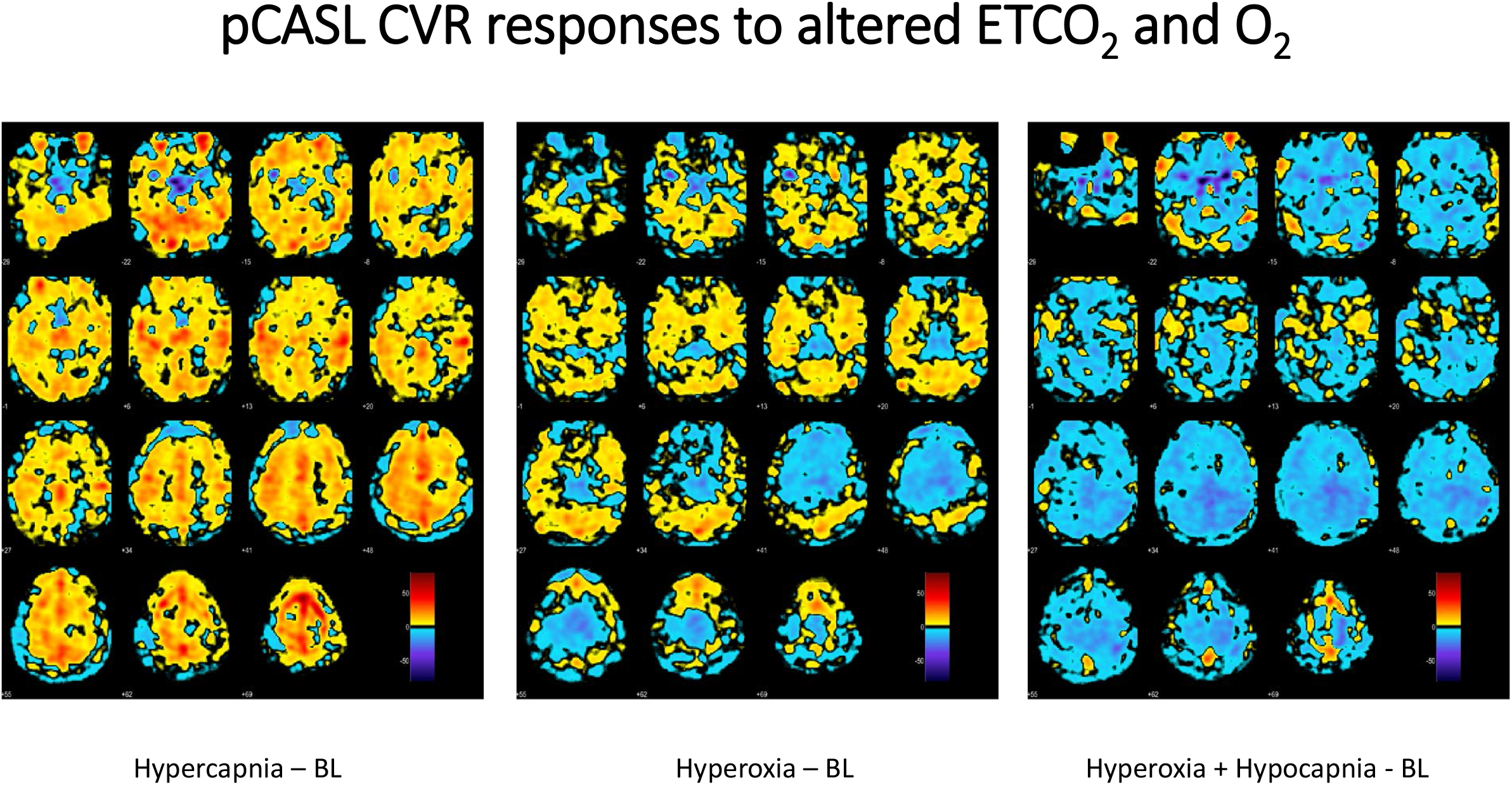
Dynamism of the response of CBF to alterations in ET gases. A: CVR difference map of hypercapnia – baseline settings at normoxia. B: CVR difference of hyperoxia – baseline settings at normocapnia, C: CVR difference of hyperoxia/hypocapnia – baseline. Such a situation is often present during anaesthesia. The large decrement in regional CBF is essentially universally evident.

**Figure 5A and B:**
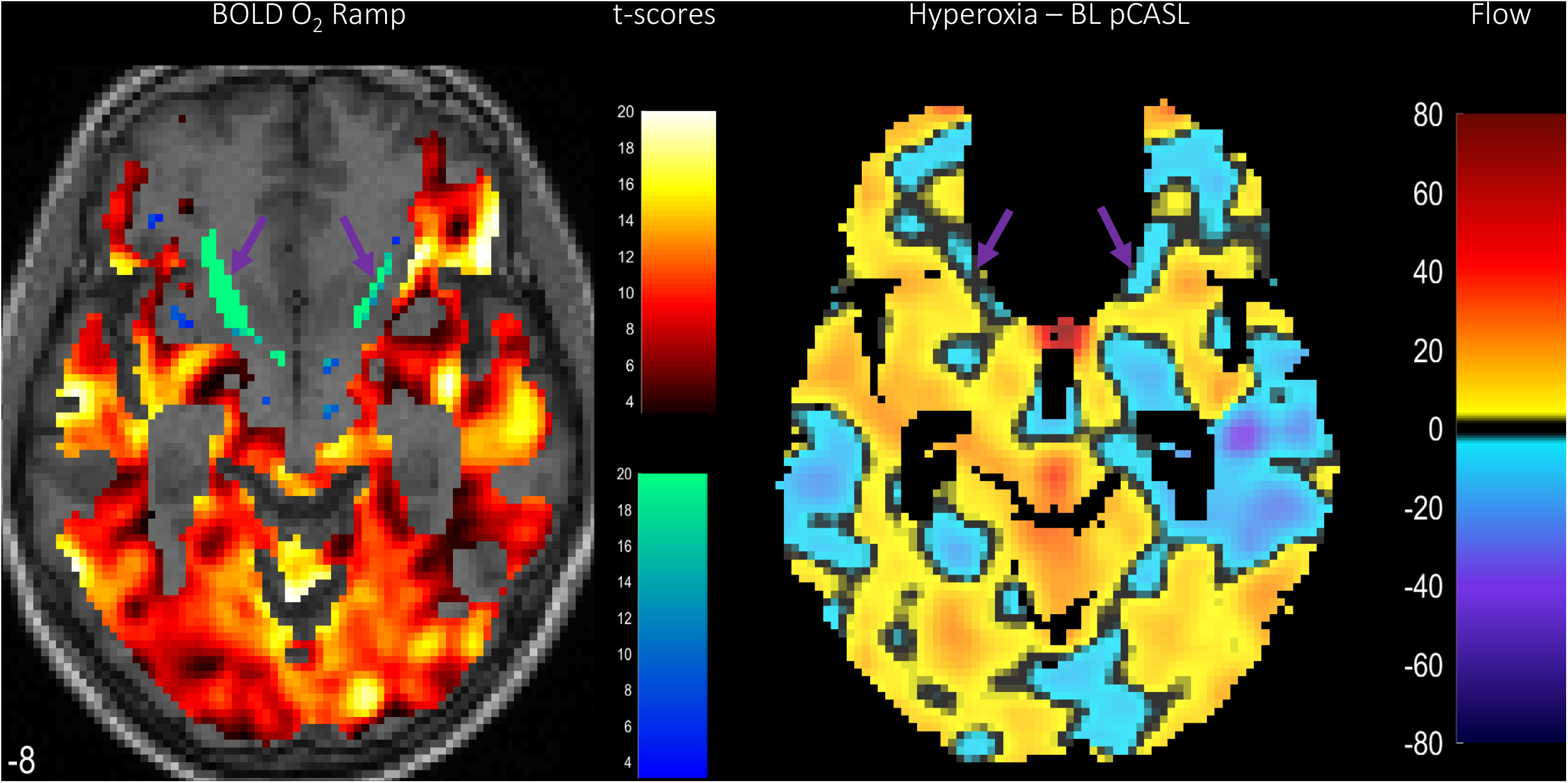
A single slice at −8 mm below the AC-PC line to show the relationship between BOLD response to hyperoxia and the changes in CBF under these conditions. The heterogeneity of response is evident as is the correlation of decreased BOLD (inverse response) and decreased CBF in regions of the frontal cortex, depicted by the purple arrows. The scales for the BOLD image are t-scores as described above. The scale for the CBF results is ± flow mg/g/min.

**Table 2.**
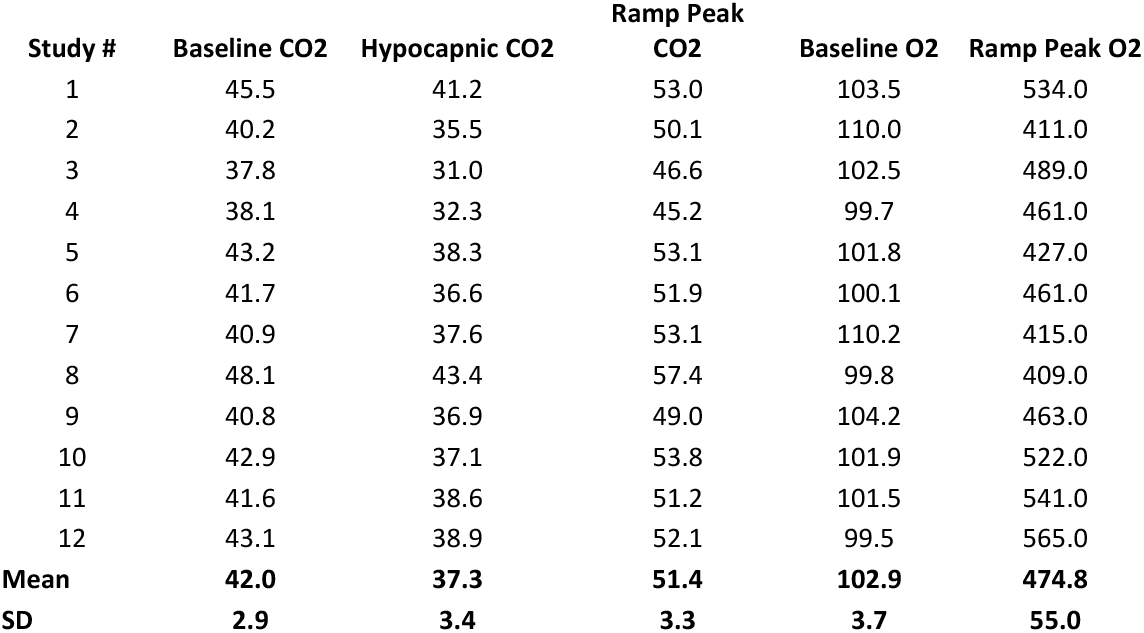
End-tidal Gas Targeting.

**Table 3.**
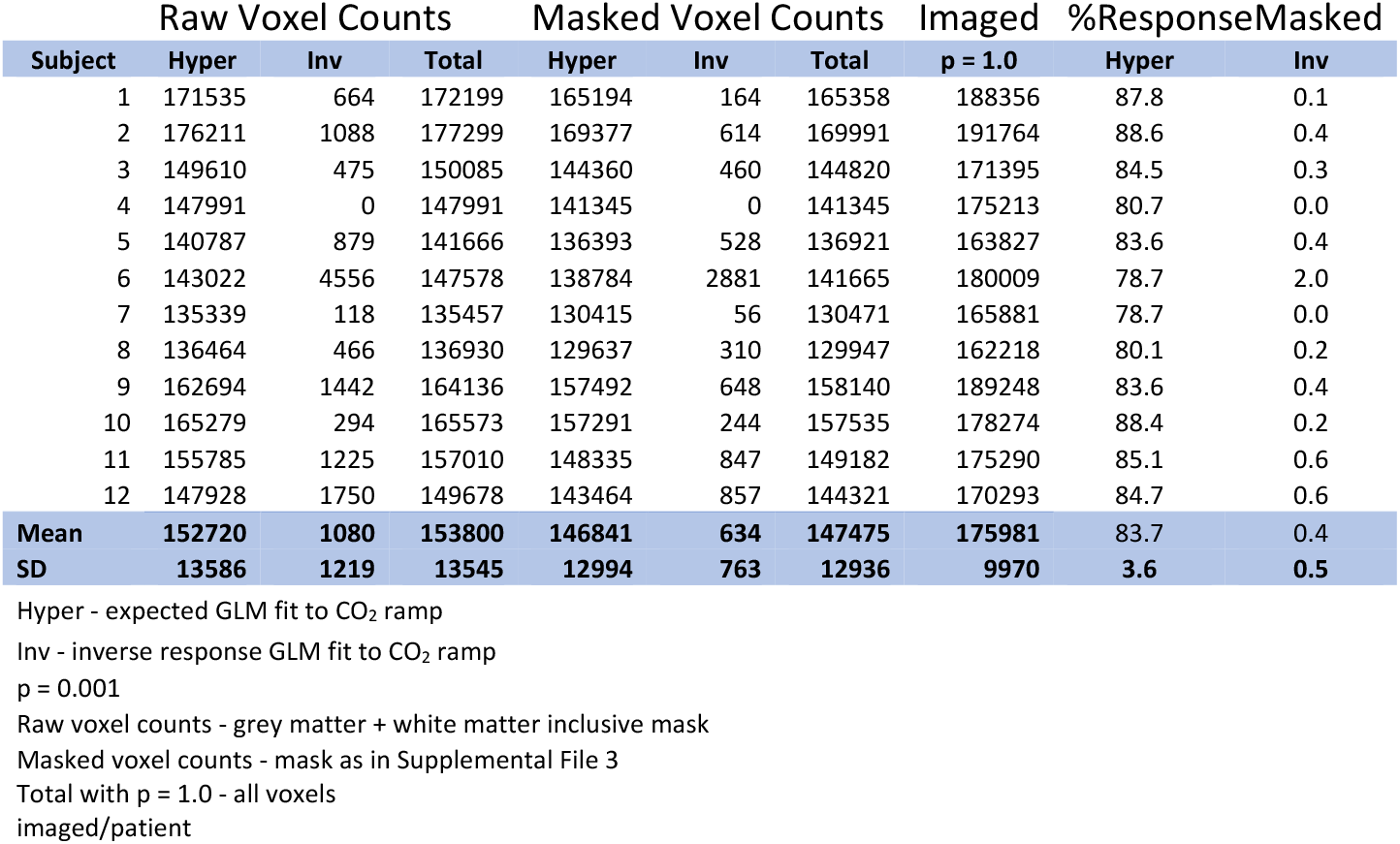
1^st^ Level Analysis CO_2_ Ramp.

**Table 4.** 1^st^ Level Analysis O_2_ Ramp.

|  | Raw Voxel Counts |  |  | Masked Voxel Counts |  |  | Imaged | %ResponseMasked |  |
| --- | --- | --- | --- | --- | --- | --- | --- | --- | --- |
| Subject | Hyper | Inv | Total | Hyper | Inv | Total | p = 1.0 | Hyper | Inv |
| 1 | 167536 | 1018 | 168554 | 161574 | 411 | 161985 | 189485 | 85.3 | 0.2 |
| 2 | 162894 | 37 | 162931 | 156581 | 0 | 156581 | 192232 | 81.5 | 0.0 |
| 3 | 104446 | 10378 | 114824 | 101379 | 8882 | 110261 | 172364 | 58.8 | 5.2 |
| 4 | 164438 | 49 | 164487 | 156126 | 41 | 156167 | 174910 | 89.3 | 0.0 |
| 5 | 158102 | 1280 | 159382 | 151166 | 882 | 152048 | 165354 | 91.4 | 0.5 |
| 6 | 161567 | 5672 | 167239 | 154975 | 3895 | 158870 | 179049 | 86.6 | 2.2 |
| 7 | 143566 | 4326 | 147892 | 138870 | 2949 | 141819 | 166975 | 83.2 | 1.8 |
| 8 | 146776 | 618 | 147394 | 138662 | 138 | 138800 | 162406 | 85.4 | 0.1 |
| 9 | 175849 | 1432 | 177281 | 168717 | 726 | 169443 | 189268 | 89.1 | 0.4 |
| 10 | 163619 | 6699 | 170318 | 155999 | 3478 | 159477 | 179612 | 86.9 | 1.9 |
| 11 | 160986 | 3395 | 164381 | 154632 | 1236 | 155868 | 175470 | 88.1 | 0.7 |
| 12 | 158532 | 1409 | 159941 | 151934 | 917 | 152851 | 170872 | 88.9 | 0.5 |
| Mean | 155693 | 3026 | 158719 | 149218 | 1963 | 151181 | 176500 | 84.5 | 1.1 |
| SD | 18258 | 3194 | 16253 | 17219 | 2567 | 15294 | 9822 | 8.6 | 1.5 |

Hyper - expected GLM fit to O<sub>2</sub> rampInv - inverse response GLM fit to O<sub>2</sub> ramp
p = 0.001
Raw voxel counts - grey matter + white matter inclusive mask
Masked voxel counts - mask as in Supplemental File 3
Total with p = 1.0 - all voxels
imaged/patient

**Table 5.**
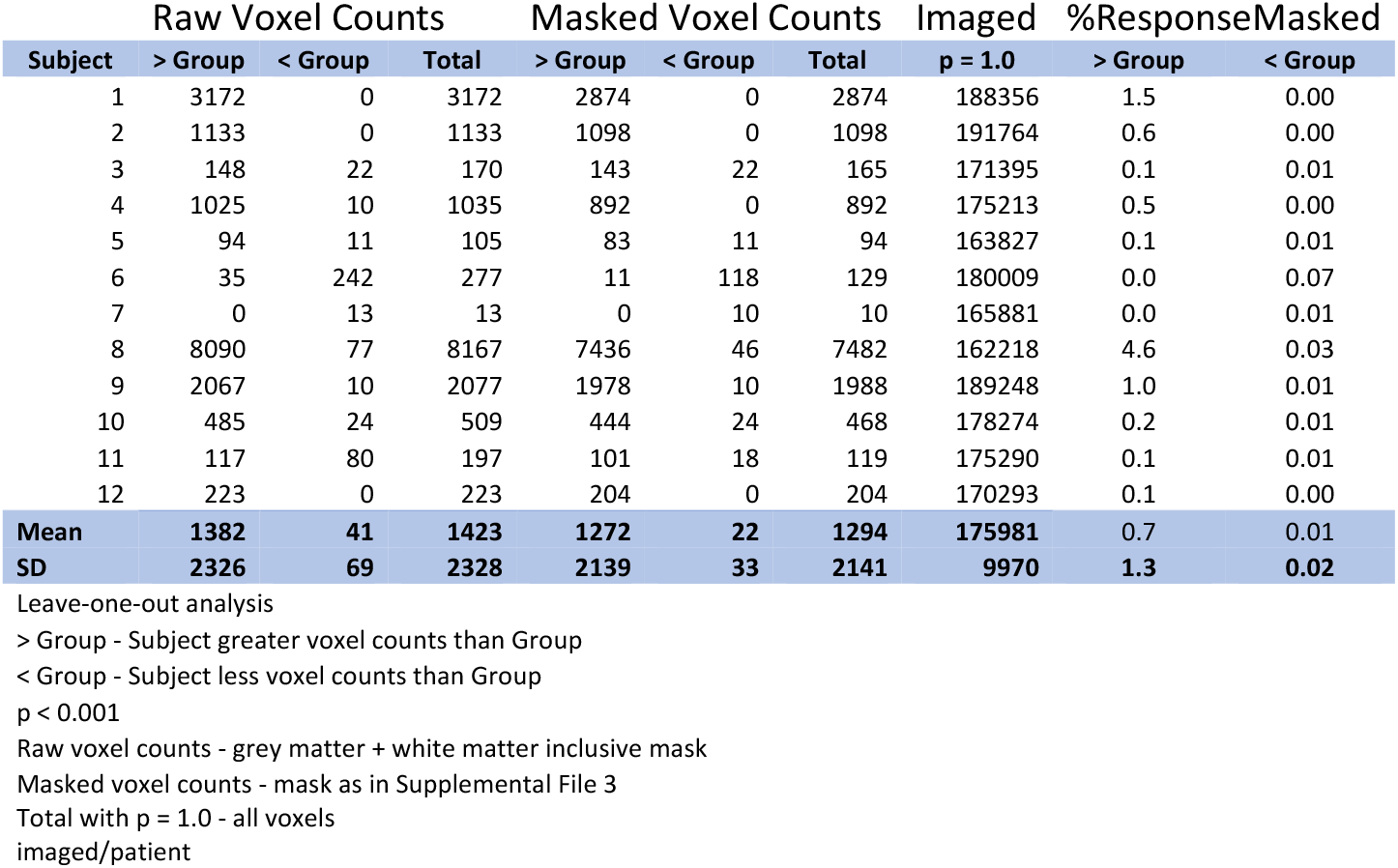
2^nd^ Level Analysis CO_2_ Ramp.

**Table 6.**
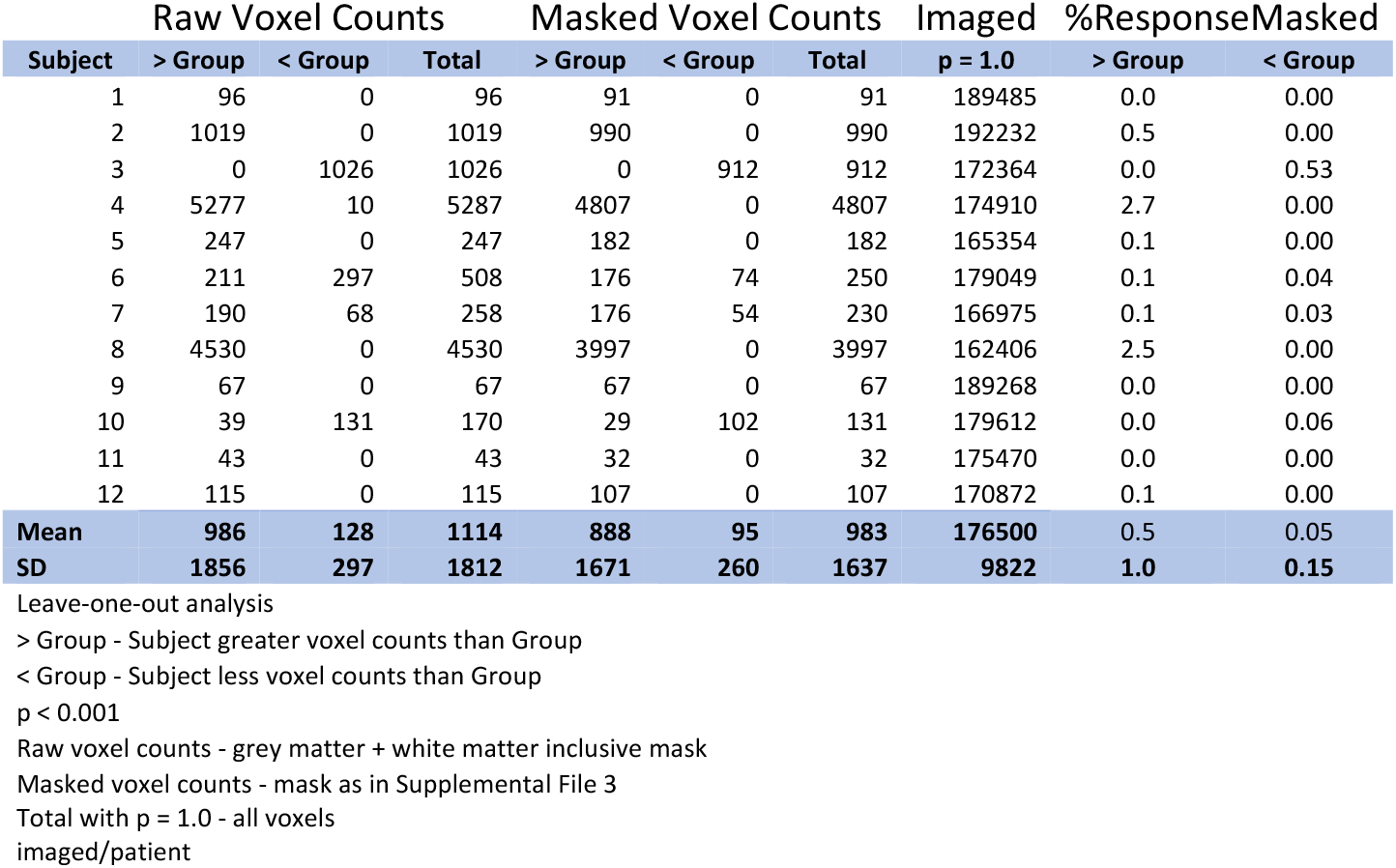
2^nd^ Level Analysis O_2_ Ramp.

## Discussion

The results of this study provide preliminary evidence of the detrimental impact of respiratory gas stress on the human brain, and have important implications for the conduct of anaesthesia on POND. This imaging study was undertaken to develop an atlas of healthy adults for comparison purposes for research protocols examining cerebrovascular reactivity (CVR) to CO_2_ and O_2_ in a number of disease states. Prior work by a number of members of this group, have used similar approaches to study concussion in adolescent patients.^29,30,31^ As these CVR studies evolved it became apparent that the findings may have considerable import regarding the conduct of anaesthesia and control of ET respiratory gases with the potential to influence the incidence and severity of perioperative neurocognitive disorder (POND). Reinforcing this is the finding that intraoperative hypocapnia is associated with an increased incidence of POND^23,32^ in adults greater than 60 years undergoing major non-cardiac surgery. Also reconfirmed is a signal indicating greater inverse CO_2_ responsiveness in older subjects (discussed in Supplemental File 4). This signal has previously been identified as a biomarker for POND in these patients after surgery.^23^ As such, examination of the potential consequences of alterations in CVR to respiratory gases during anaesthesia and surgery appears warranted.

Anaesthetic management comprises more than just administration of controlled doses of either volatile or intravenous agents to render the patient insensate. Intraoperative management also includes haemodynamic control and breath-to-breath management and monitoring of respiratory gases – O_2_ and CO_2_. Oxygen is usually given in increased fractional concentrations over that seen in room air (FiO_2_ = 0.21). It is not uncommon to administer a significantly elevated O_2_ (such as an FiO_2_ = 0.50) for the duration of a surgical case and FiO_2_ = 1.0 is virtually always given at the start and end of surgical cases as a buffer for the risk of hypoxia. Except for short surgical procedures, patients are usually mechanically ventilated which is frequently associated with lower tensions of ET CO_2_ for the duration of the intraoperative procedure.^33,34^ High tensions of CO_2_ are also not uncommon at the end of a procedure with reversal of muscle relaxants and reestablishment of spontaneous ventilation. Thus, even during a routine anaesthetic, respiratory gases usually have a wide range of fluctuation (far outside the range seen in the awake healthy state). The potential for large variation in CBF with such alterations in ET gases during conduct of even a normal anaesthetic can be deduced by examination of the various images from this study. A marked dynamism is clear and in certain subjects ‘steal’ phenomena are seen where the inverse response to expected changes in BOLD signal or CBF to CO_2_ alterations and O_2_ are evident. This is especially pronounced in the frontal cortex where executive function resides, potentially placing this critical area of the brain at risk with alterations in respiratory gases during anaesthesia.

Mechanistically the inverse responses to changing ET tensions of CO_2_ and O_2_ differ. The ‘intracranial steal’ seen with hypercapnia is well elucidated.^35^ Regional areas of vasoparalysis result in shunting of blood from their perfused areas to adjacent regions with increasing adjacent regional vasodilation with hypercapnia. The regional ‘intracranial steal’ seen with hyperoxia appears the reverse. Excessive regional vasoconstriction seems evident. Surrounding areas with less vasoconstriction receive higher flow. The inverse BOLD response suggests vasoconstriction of such intensity that venular offloading of oxygen from hemoglobin is excessive here, resulting in increased paramagnetic signal.^36,37^ This explanation suggests excessive regional vascular tone in these feeding vessels – the mechanism to be further elucidated. It has been noted however, that frontal neurons in Alzheimer transgenic mice are vulnerable to hyperoxia through synaptic dysfunction and brain oxidative stress.^38^ Vasoconstriction with hyperoxia is a proposed mechanism in this model.^39,40^ Examination of the inverse BOLD responses seen in the BOLD signal curve fits in Figure 1 and 2 indicates that both the response to CO_2_ and O_2_ alterations are rapid and incremental, indicating a very dynamic process. The pCASL findings indicate that the regional vasoconstrictive influence of hyperoxia is amplified in the presence of hypocapnia (Figure 4). As noted above, this combination of ET gas conditions is ubiquitous during the conduct of anaesthesia. The curve fit examining the magnitude of inverse voxels fits a second order polynomial correlated to increasing age although the sample size is small. The best fit to a paraboloid, however, is an interesting finding as the incidence of POND is bimodal at the extremes of life. Whether inverse voxel signatures are more evident in neonates and young children is an open question.

These control studies have been conducted in awake individuals so the impact of anaesthesia – both from volatile agents (known cerebral vasodilators) and intravenous agents (know cerebral vasoconstrictors with the exception of ketamine) may alter the response to CO_2_ and O_2_ as seen here. However, a carefully done study in rats anaesthetized with isofluane (a volatile agent) reports many of the same BOLD and CBF findings using similar imaging approaches, suggesting that the respiratory gas effects on cerebral perfusion translate to the anaesthetized state with volatile agents.^41^ Recent work by Venkatraghavan et al.^42^ showing marked fluctuations in brain BOLD in 4 patients undergoing propofol anaesthesia with intracranial steal using a similar CO_2_ ramp protocol as in this study, indicating that the CO_2_ responsiveness we describe in the awake state persists under anaesthesia. Also well-known is the rapid response by the brain to alterations in ET CO_2_ to control intracranial volume during anaesthesia for neurosurgical procedures. We have previously demonstrated that intraoperative hypocapnia is associated with an increased incidence of POND,^23^ however, ET O_2_ fluctuations were not considered as a potential factor in that study. The results of the present study suggest both O_2_ and CO_2_ changes may be important. MacDonald et al.^43^ reported variations in CBF of nearly 100% in 10 healthy subjects exposed to hyperoxia. This heterogeneity of regional responses to hyperoxia suggests that one CBF does not fit all regions, either in grey or white matter. Thus, each individual may have a unique CBF response during surgery with great regional heterogeneity as the ET gases are manipulated. Other human studies have indicated that hyperoxia induces cerebral vasoconstriction as a consequence of altered regional nitric oxide potentially contributing to the stress diathesis under anaesthesia.^44^ Also of note are findings of altered CVR with increasing age^45^ and with signs of dementia in the elderly^46^ or in patients with biomarkers such as APO4ε related to early onset Alzheimer’s disease.^47^ The identification of regional ‘intracranial steal’ or ‘blue brain’ on the BOLD images with a CO_2_ ramp protocol is increasingly being used to establish the need for and measure results from revascularization with superficial temporal artery to middle cerebral artery (STA-MCA) anastomoses in patients with severe intracranial vascular compromise.^48,49^ Such patients when successfully revascularised see improvements in CVR and complete or partial resolution of the ‘blue brain’ signal that indicated the area of regional steal. Thus, these abnormal regions as identified in our study have clear pathological correlates. There are currently no comparable studies regarding O_2_ ramp protocols and the ‘blue brain’ seen in this study with this imaging protocol but analogies to the findings with the CO_2_ ramp protocol are envisioned. Also, importantly not addressed here is the potential findings of an O_2_ ramp protocol in the presence of a clamped period of hypocapnia. It is this sequence that is especially interesting as this respiratory gas profile is common during anaesthesia. The short hypocapnic periods obtained in this study were obtained by having the subject spontaneously hyperventilating on command. An 11-minute ramp sequence with stable hypocapnia by spontaneous hyperventilation for such a duration is not feasible in awake subjects. That said our 3^rd^ pCASL sequence in 4 subjects where a 3-minute period of hyperoxia with hypocapnia indicates a synergic diffuse decrease in CBF which potentially could have deleterious effects in patients at risk of POND.

Based on these initial observations we advance the hypothesis that end-tidal respiratory gas control is a critical management consideration during anaesthesia; a much more important consideration than previously thought. Older patients appear to be at greater risk of ‘intracranial steal’ of regional CBF with alterations in CO_2_ and O_2_. Our previous study confirms an association between CO_2_ management and POND.^23,32^ The imaging in the present study suggests that a combination of hyperoxia and hypocapnia may be particularly worrisome. We suggest that anaesthetic management may be optimized for cerebral health by attempting to maintain respiratory gases at or near the patient baseline values to stabilize CBF during the conduct of anaesthesia. An exception may occur during neurosurgical procedures where deliberate alterations to lower ET CO_2_ may be required. The marked CBF fluctuations as seen with controlled ET gas manipulation in this study imply that such CBF changes can occur under anaesthesia where similar changes in ET gases routinely occur. Such large alterations in CBF and oxygenation may be a contributing factor to the development of POND in susceptible individuals. A clinical trial comparing rigorous maintenance of end-tidal gases at a patient’s own baseline, to standard anaesthetic management to determine if the changes in CBF and oxygenation as seen in awake subjects in this study have an influence on the development of POND could be considered.

## Acknowledgements

The authors thank Dr. Joseph Fisher, Dr. James Duffin and Dr. David Mikulis, developers of the RespirAct used in these studies for helpful discussions about the methodological approach used here for ET gas control.

## Authors Contributions

WACM: Funding, conception of the study, recruitment, conduct of the study, data collation, data processing, analysis, writing

REl-G: Funding, conception of the study, data collation, data processing, analysis, writing

LR: Conception of the study, data collation, data processing, analysis, writing

JP: Recruitment, conduct of the study, data collation, data processing, analysis, writing

ME: Funding, conception of the study, analysis, writing

KK: Recruitment, conduct of the study, data collation, data processing, writing

KF: Recruitment, conduct of the study, data collation, data processing, writing

MRG: Funding, conception of the study, data collation, data processing, analysis, writing

---

**Supplemental File 1:**
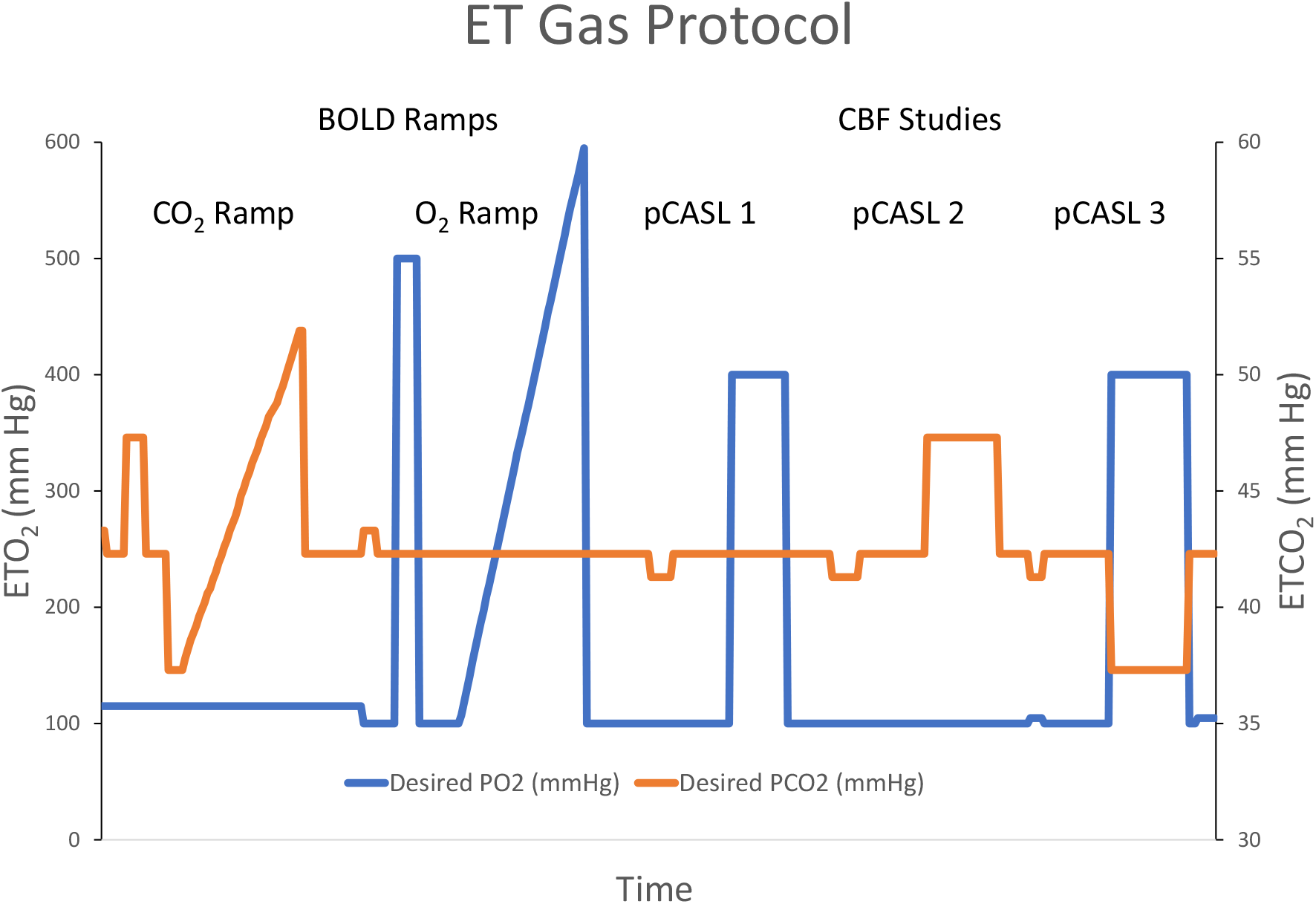
Display of the end-tidal gas control as described in Methods for one subject.

**Supplemental File 2:** Description of the MRI sequences used in the study.

Images were acquired using a Siemens Verio 3.0 T MR scanner with a 12-channel phased-array head coil. Anatomical imaging was acquired without manipulation of end-tidal gases using a sagittal 3D T1 MPRAGE (whole brain coverage; matrix: 256 × 256; slice thickness: 2.2 mm; no interslice gap; voxel size 2 mm × 2 mm × 2 mm) and axial gradient recalled echo planar (GRE) sequences to screen for cerebral microhemorrhages as well as GRE B0-field mapping. CVR was assessed using continuous BOLD MRI during MPET CO_2_ targeting. BOLD MRI data were acquired using a T2*-weighted single-shot gradient echo pulse sequence with echoplanar readout (field of view: 24 cm × 24 cm; matrix: 64 × 64; TR: 2,000 ms; TE: 30 ms; flip angle: 85°; slice thickness: 5.0 mm; interslice gap: 2.0 mm; voxel size 3.75 × 3.75 × 6 mm; number of temporal frames = 330; 10 s of initial imaging data was discarded to allow for equilibration). Global mean resting CBF was assessed using pseudo-continuous arterial spin labeling (pCASL) that included an initial M0 scan—Siemens ep2d_pCASL—echo planar readout (field of view 24 cm × 24 cm, TR 8,000 ms, TE 12 ms, contrast with a flip angle 90°, 20 slices, CASL method—multislice, label offset 90 mm, post label delay 1,200 ms, crusher gradient 0 s/mm^2^, and voxel size 3.8 mm × 3.8 mm × 5.0 mm). The formal pCASL sequence then followed consisting of an echo planar readout (field of view 24 cm × 24 cm, TR 4,000 ms, TE 12 ms, contrast with a flip angle 90°, 20 slices, slice thickness 5.0 mm, CASL method— multislice, label offset 90 mm, post label delay 1,200 ms, crusher gradient 0 s/mm^2^; voxel size 3.8 mm × 3.8 mm × 5.0 mm). Imaging duration was for 3 min. The first two labeled–non-labeled pairs were discarded.

**Supplemental File 3:**
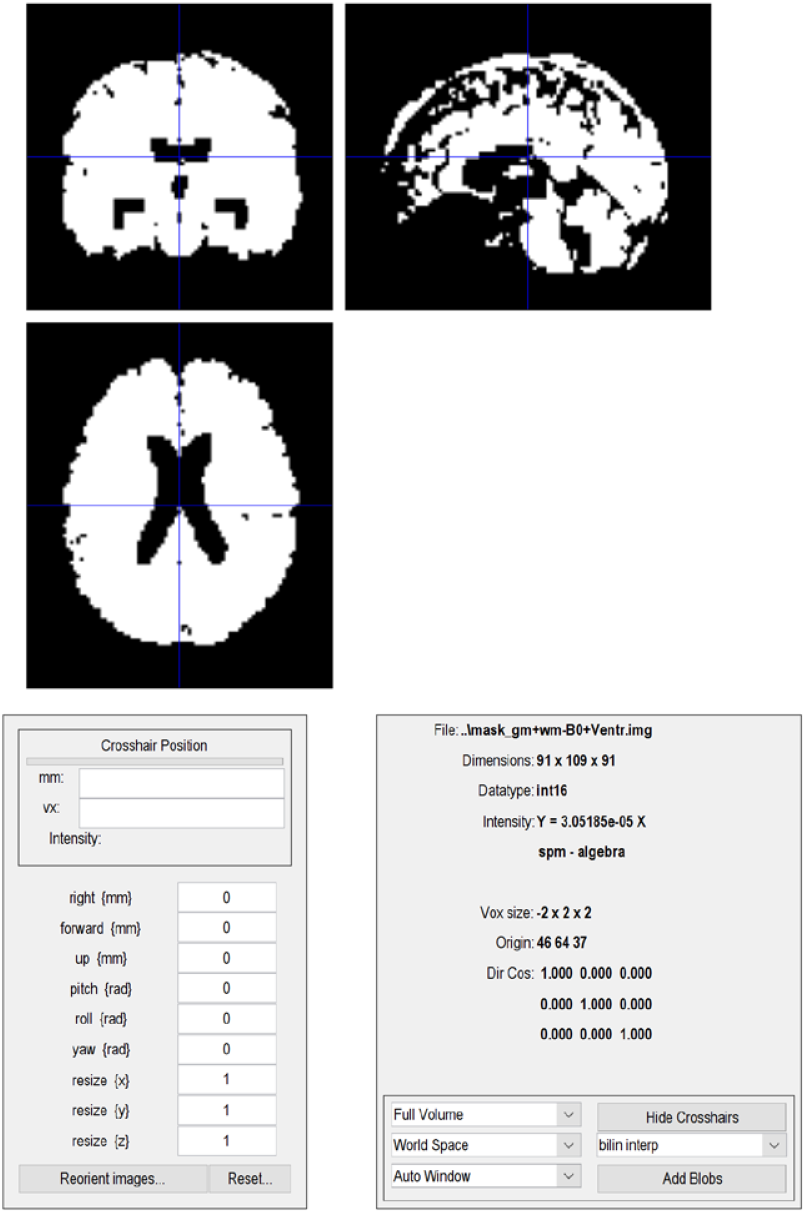
Single subject mask (individualized based on their imaging) with B0 inhomogeneities masked out and a CSF ventricular mask included at 1-fold dilation as determined from the WFU pick atlas. In this manner potential BOLD signal error were removed from areas adjacent to surface transitions were not included in the voxel analysis at the 1^st^ and 2^nd^ levels.

**Supplemental File 4:**
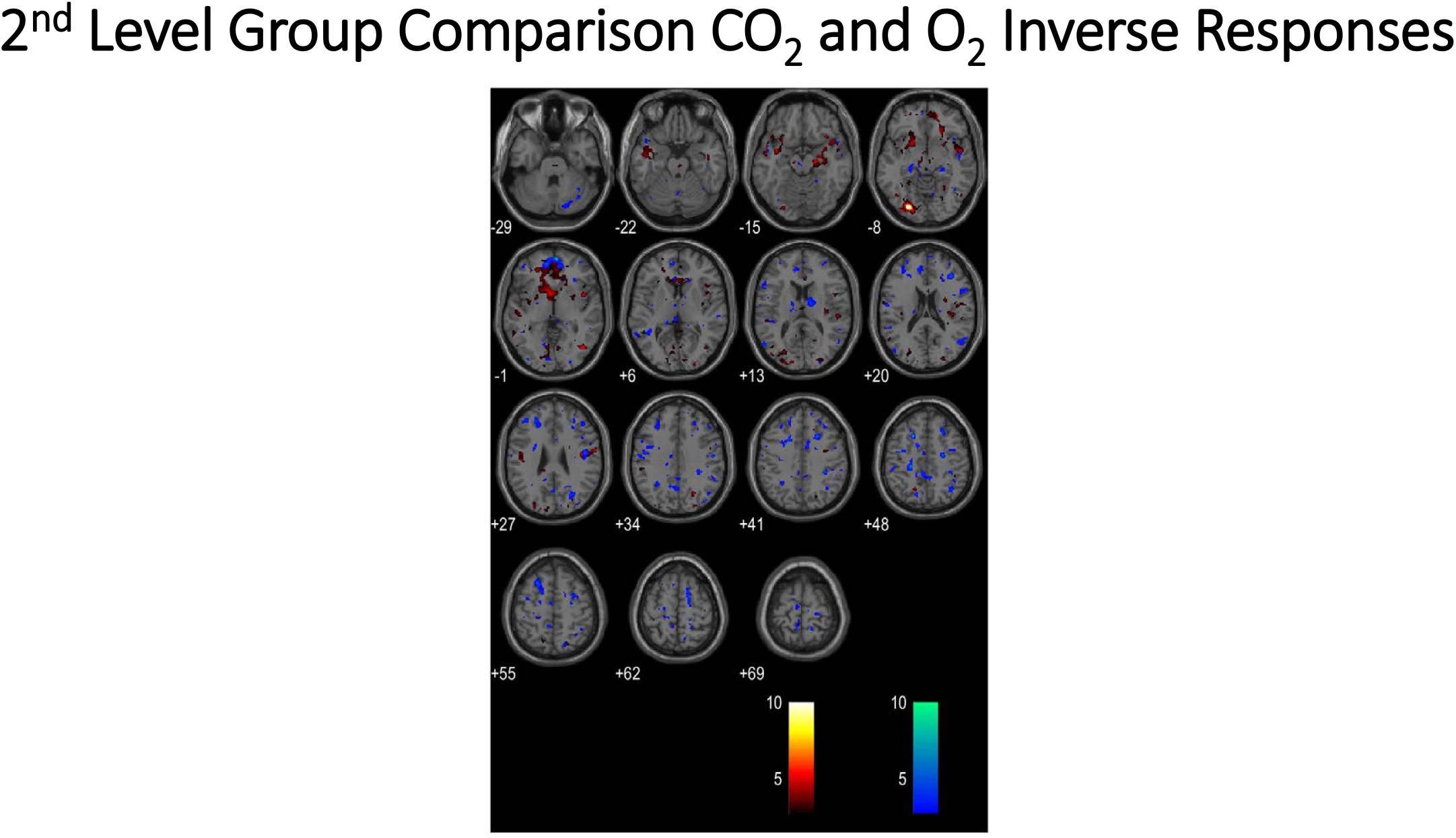
2^nd^ level group analysis based on the inverse response to the O_2_ ramp (orange voxels) and the inverse response to the CO_2_ ramp (blue voxels) at the p = 0.01 level. The groups were 4 older subjects age (53-62) with voxel counts above 50 at the p = 0.001 level from Table 5 and 6. The ‘control group’ was the other 8 subjects (mean age 43 ± 13). The distribution of blue voxels showing the group response of a greater voxel count to the inverse CO_2_ stimulus is similar to that seen in reference 23 and identified as a biomarker of POND in this previous work.

